# Assessing epidemiological parameters and dissemination characteristics of the 2000 and 2001 foot-and-mouth disease outbreaks in Rio Grande do Sul, Brazil

**DOI:** 10.1101/2022.05.22.492961

**Authors:** João Marcos Nacif da Costa, Luis Gustavo Corbellini, Nicolás Céspedes Cárdenas, Fernando Henrique Sauter Groff, Gustavo Machado

## Abstract

Since 1998 the state of Rio Grande do Sul, Brazil, has been free of foot-and-mouth disease (FMD) with yearly mandatory vaccination, until the 2000 and 2001 reintroductions. This study gathers data from both outbreaks including official veterinary state service archives and field investigation reports to quantify epidemiological parameters such as epidemic duration, number of secondary infected farms and animals, and estimate the epidemic rate of growth. We apply a Bayesian latent variable approach to estimate the time-varying reproduction number and calculate new confirmed cases by infection date. Additionally, we utilized between-farm animal movements to reconstruct possible FMD transmission and characteristics of spread over the current at-risk population, by incorporating bovine movement data from 2018 to 2020 as standard to benchmark infected network parameters. The results were consistent with the reports generated by the official investigation of the outbreaks and the models and results presented in this study may be useful for assessing the transmission dynamics and support control measures in the future.

## 1. Introduction

Foot-and-mouth disease (FMD) affects a wide variety of cloven-hoofed domestic and wild animals, although pathogenesis has been studied mainly in cattle and pigs (Arzt, Baxt, et al., 2011; Grubman & Baxt, 2004; Kitching & Alexandersen, 2002). The phases of FMD are characterized by a pre-symptomatic infectious period followed by sustained viremia accompanied by the onset of the symptoms, and a convalescence period with the resolution of clinical disease (Arzt, Baxt, et al., 2011; Arzt, Juleff, et al., 2011; Grubman & Baxt, 2004; Kitching & Alexandersen, 2002; Musser, 2004). Direct contact between infected and susceptible animals is the main transmission route because the FMD virus is often present in the body fluids of acutely infected animals and aerosol shedding. The indirect exposure to the excretions and secretions of acutely infected animals or uncooked meat products may be a source of infection to susceptible animals (Colenutt et al., 2020; Mielke & Garabed, 2020; Paton et al., 2018; Rueda et al., 2015). FMD control and eradication are challenging and expensive, mainly because of international trade bans and the direct and indirect costs of control and eradication activities (Junker et al., 2009; Kitching & Alexandersen, 2002; Pendell et al., 2007). The estimated annual FMD cost in countries with endemic areas ranges from US$ 6.5 to US$ 21 billion, and the economic losses by the occurrence of cases in free areas can exceed US$ 1.5 billion per year (Knight-Jones & Rushton, 2013). In the United States, the reintroduction of FMD was estimated to cost approximately US$47 billion in gross domestic product loss and 677,000 jobs lost would be a consequence of a depopulation strategy with no use of vaccination (Miller et al., 2019).

In 2000, the reintroduction of FMD into part of South America caused extensive economic and social losses to the livestock production chain, in Argentina, Uruguay and the State of Rio Grande do Sul, Brazil (Lyra & Silva, 2004; MAPA, 2002). The areas in Brazil territory were recognized as “free with vaccination” by OIE just two years before. The rapid spread of FMD to a region considered free of FMD led to an increase in prevention, surveillance, and eradication strategies levels, and the virus was considered to have been eliminated (Lyra & Silva, 2004; Mayen, 2003). However, in 2001, a rapid and massive epidemic rapidly affected the Prata basin region (Correa Melo et al., 2002; Lyra & Silva, 2004; Naranjo & Cosivi, 2013; Perez et al., 2004a). Since 2001 the number of outbreaks in South America has decreased significantly (Naranjo & Cosivi, 2013). This trend was interrupted by the FMD type O virus events in Colombia in 2017 and 2018 (FAO, 2018; Gomez et al., 2019). Since 2019 no new outbreaks have been reported in the region, despite Venezuela’s absence of official international status for FMD (PANAFTOSA, 2020). In 2021, Rio Grande do Sul, among other Brazilian states, was recognized by OIE as free from FMD without vaccination (OIE, 2021).

In order to achieve and maintain recognition of animal health status in a region, strategic policies must be based on knowledge about the characteristics of the disease agent as well as the disease dynamics in the livestock populations, making it possible to establish interventions (Gonçalves & de Moraes, 2017; Paton et al., 2018). Mathematical and simulation models may be applied to evaluate the effectiveness of policies to mitigate the impact of epidemics and guide the routine programs (Knight-Jones et al., 2016; Pomeroy et al., 2017; Probert et al., 2018). There is an urgent demand to estimate transmission parameters directly of outbreaks, which may also include farm-level information, animal populations, husbandry characteristics, and movement patterns (van Andel et al., 2021).

The reproduction number has been largely used to estimate the transmission potential of FMD during outbreaks (Arjkumpa et al., 2021; Colenutt et al., 2020; Estrada et al., 2008; Ferguson et al., 2001; Keeling, 2005; Muroga et al., 2012; Perez et al., 2004a; Tadesse et al., 2019). The basic reproduction number (*R*_0_), is often used to estimate the spreading capacity of an infectious agent newly introduced to a totally susceptible population, briefly, it represents the number of secondary infections caused by a single case in that population (Anderson & May, 1995; Delamater et al., 2019; Dietz, 1993). As epidemics continue to propagate, numerous factors can directly influence transmission dynamics, such as the availability of susceptible animals, the effectiveness of control measures carried on by Animal Health Services, and spatial correlations between farms, demanding an estimate that accounts for the changes in the disease spread pattern (Haydon et al., 2003; Tildesley & Keeling, 2009). Thus, for ongoing epidemics, the appropriate methodology would be the estimation of the time-varying reproduction number (*R*_*t*_), which is defined as the average number of secondary cases per primary case at a given time *t* (Cori et al., 2013; Merl et al., 2009; Vegvari et al., 2021). To reduce distortions on the epidemic curve in outbreaks where the case counting is obtained by the date of notification the use of models that accounts for uncertainty, associated with delays between symptoms onset and the date of notification, would be an appropriate approach (Abbott et al., 2020; Gostic et al., 2020; Nakajo & Nishiura, 2022; Probert et al., 2018).

Other useful models to estimate transmission dynamics and epidemiological parameters are based on the retrospective reconstruction of previous outbreak networks (Firestone et al., 2020; Hayama et al., 2019; Jombart et al., 2014). Such network modeling exercises emulate the characteristics of spread and allow the identification of potential super spreading events as well as epidemics sizes by describing interactions between individuals using animal movement data (Cabezas et al., 2021; Cardenas et al., 2021, 2022; Dubé et al., 2009; Fèvre et al., 2006; Machado et al., 2021; Ruget et al., 2021; VanderWaal et al., 2016).

The objective of this study is to describe and estimate transmission parameters from the 2000 and 2001 FMD outbreaks in the State of Rio Grande do Sul. Here we gather outbreak data of outbreaks to quantify epidemiological parameters such as epidemic duration, number of secondary infected farms and animals and estimate the epidemic rate of growth. We apply a Bayesian latent variable approach using back-calculation to estimate the time-varying reproduction number and calculate new confirmed cases by infection date. Additionally, we utilized between-farm animal movements to reconstruct possible FMD transmission and characteristics of spread over the current at-risk population. Ultimately, this study provides a comprehensive understanding of historical events and supports strategic policies based on further transmission models that can be used to investigate future epidemics and the effectiveness of countermeasure actions.

## 2. Material and methods

### 2.1 FMD outbreak data

In this study we utilized 2000 and 2001 FMD epidemic data available from the state of Rio Grande do Sul official veterinary services archives (SEAPDR-RS, 2021). The databases contained information on field investigation reports produced during and after outbreaks (MAPA, 2002), including herd sizes, species, geolocation, the chronology of control and eradication events (e.g notifications, movement restrictions, and vaccination), clinical investigation findings, which included ages of clinical lesions (EUFMD, 2020; Kitching & Alexandersen, 2002; MAPA, 2002), and the number of daily new infected herds along with the population in each farm. Additionally, we also used between-farm cattle movement records from 2018 to 2020, which included the number of animals transported and the date of each shipment, as a standard to benchmark the transmissions between farms in the 2000 and 2001 outbreaks. The movement data was also provided by the official veterinary service under a data use agreement with the Rio Grande do Sul Secretary of Agriculture, Livestock Rural Development in Brazil (SEAPDR-RS, 2021).

### 2.2 Estimating FMD effective reproduction number

We implemented a Bayesian latent variable approach to calculate the time-varying reproduction number using daily infected animals based on the notification dates between August 1^st^ to September 22^nd^, 2000, and from May 5^th^ to July 18^th^, 2001. We reconstructed the outbreak time series to estimate the latent infections *I*_*t*_ by fitting a back-calculation model. The infection estimates were then mapped to a mean reported case count *D*_*t*_ given an incubation period and reporting delay distributions convolved into *ξ*. Observed reported cases *C*_*t*_ were finally generated from a negative binomial model with mean *D*_*t*_ and overdispersion *ϕ* from an exponential prior with mean 1. Finally, the time-varying reproduction number *R*_*t*_ was estimated by the ratio of the number of new infections generated at time step *t*, to the sum of infection incidence up to time *t-1* weighted by an uncertain generation time function *W*_*s*_ (Abbott et al., 2020, 2021; Cori et al., 2013; Sherratt et al., 2021).

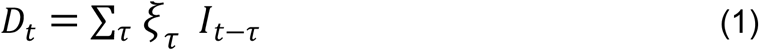

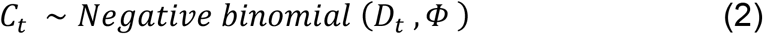

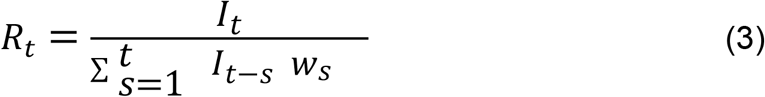

The generation time with a mean of 6.1 and standard deviation of 4.6 was obtained from the literature (Haydon et al., 2003). For the incubation period, mean and standard deviation were drawn from a Poisson distribution with λ equal to 5.9 based on a published meta-analysis (Mardones et al., 2010). The delays between symptom onset and case reporting for each infected farm were extracted directly from the 2001 FMD outbreak investigation reports (SEAPDR-RS, 2021). Then, we fitted a log-normal distribution to a subsampled bootstrap of the delay data. The Bayesian latent variable model was implemented in R version 4.1.1 using the EpiNow2 package (Abbott et al., 2021; R Core Team, 2019). For every model run, four chains were used with a warmup of 500 samples each and 4,000 samples post-warmup.

### 2.3 Static network analysis

The contact network uses between-farm cattle movement data from January 2018 to December 2020, as described earlier. Data regarding the location and official registry of infected farms in 2000 and 2001 outbreaks were assessed and validated in order to correspond with current movement data. We constructed contact networks in which farms were defined as nodes and movements between farms were considered as unweighted and directed edges (Wasserman & Faust, 1994). In order to reconstruct the infected networks, we extracted two subsets of the full contact network, in which the group of farms involved in the 2000 and the group of farms involved in the 2001 outbreak were assigned as infected independently. The infected network (g) was represented as a directed graph and comparisons were made using the following parameters: the number of nodes, edges, fragmentation, a total of animals moved, mean of the shortest paths length, and the estimates for network parameters graph density, giant strongly connected component (GSCC), and giant weakly connected component (GWCC) (Dubé et al., 2009; Wasserman & Faust, 1994). We also use the k-test, a permutation-based approach, to characterize the pattern of the infected nodes within the contact network and to indicate if the cattle movement would explain transmission pathways between farms (VanderWaal et al., 2016). The observed distribution of the outbreak farms in the full static network was compared independently for each outbreak year, with the expectation under the null hypothesis of random distribution of cases (i.e. infected nodes) within the network. The k-statistic is the mean number of cases observed within one step of an infected node in the network. The null distribution was generated through 1000 Monte Carlo simulations in which the cases in the population were randomly redistributed in the network. Analyses were conducted in R version 4.1.1 using the igraph package (Csardi, G & Nepusz, T, 2006; R Core Team, 2019).

## 3. Results

The reintroduction of FMD in the state of Rio Grande do Sul in 2000 and 2001 were considered independent events. The first official notification was made on August 1^st^, 2000, at the municipality of Jóia, 145 km from the border with Argentina (Figure 1). The outbreak spread into three other neighboring municipalities, Eugênio de Castro, Augusto Pestana, and São Miguel das Missões, in the northwest region of the state corresponding to a total area of 3,439 km^**2**^. A total of 22 farms (Table 1) were infected and the outbreak was confirmed to be caused by the type O virus (MAPA, 2002).

**Figure 1.**
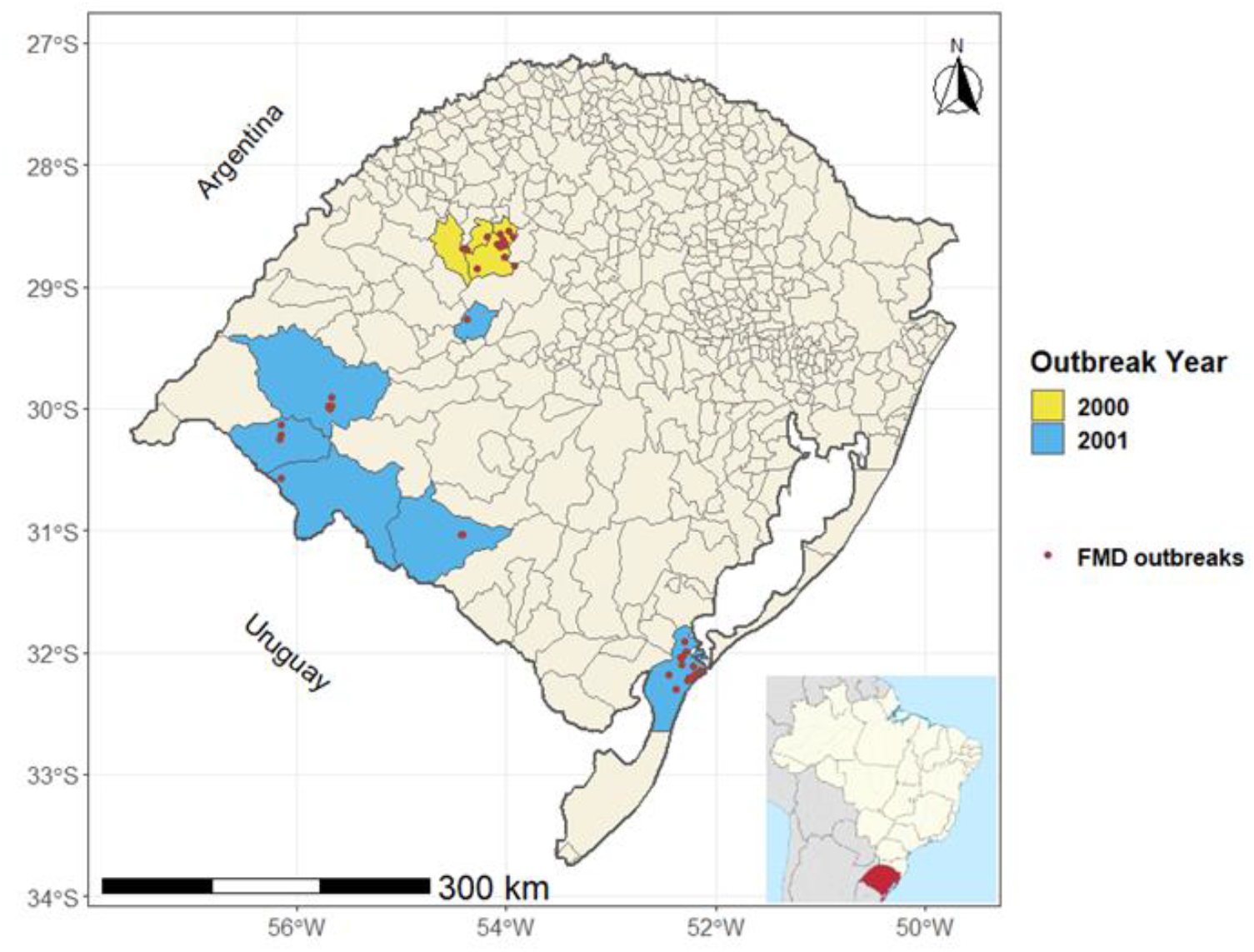
The distribution of two FMD outbreaks in Rio Grande do Sul, Brazil. Municipalities involved in the outbreaks are represented by yellow (2000) and blue (2001) areas. Infected farms’ locations are represented by red dots.

**Table 1.**
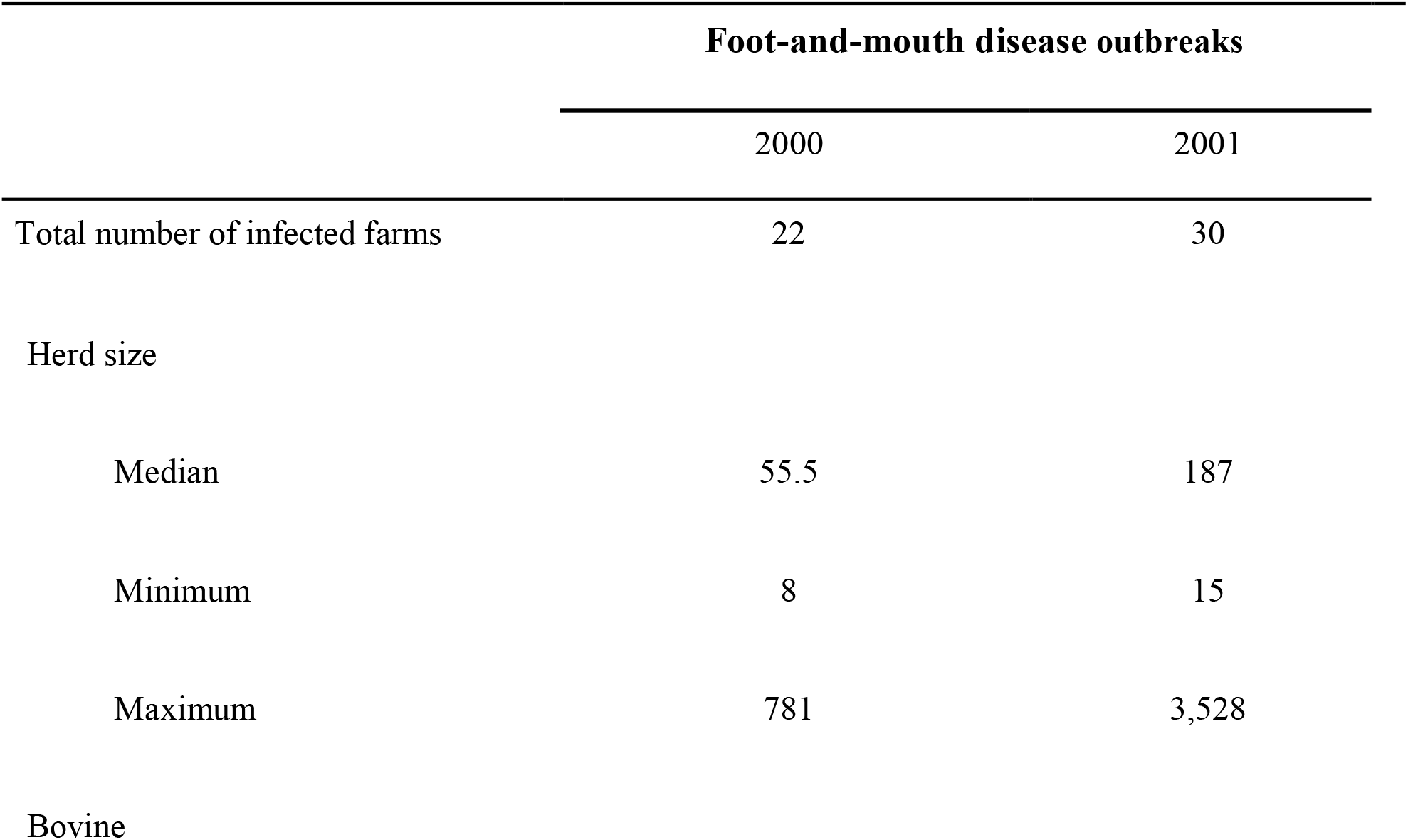
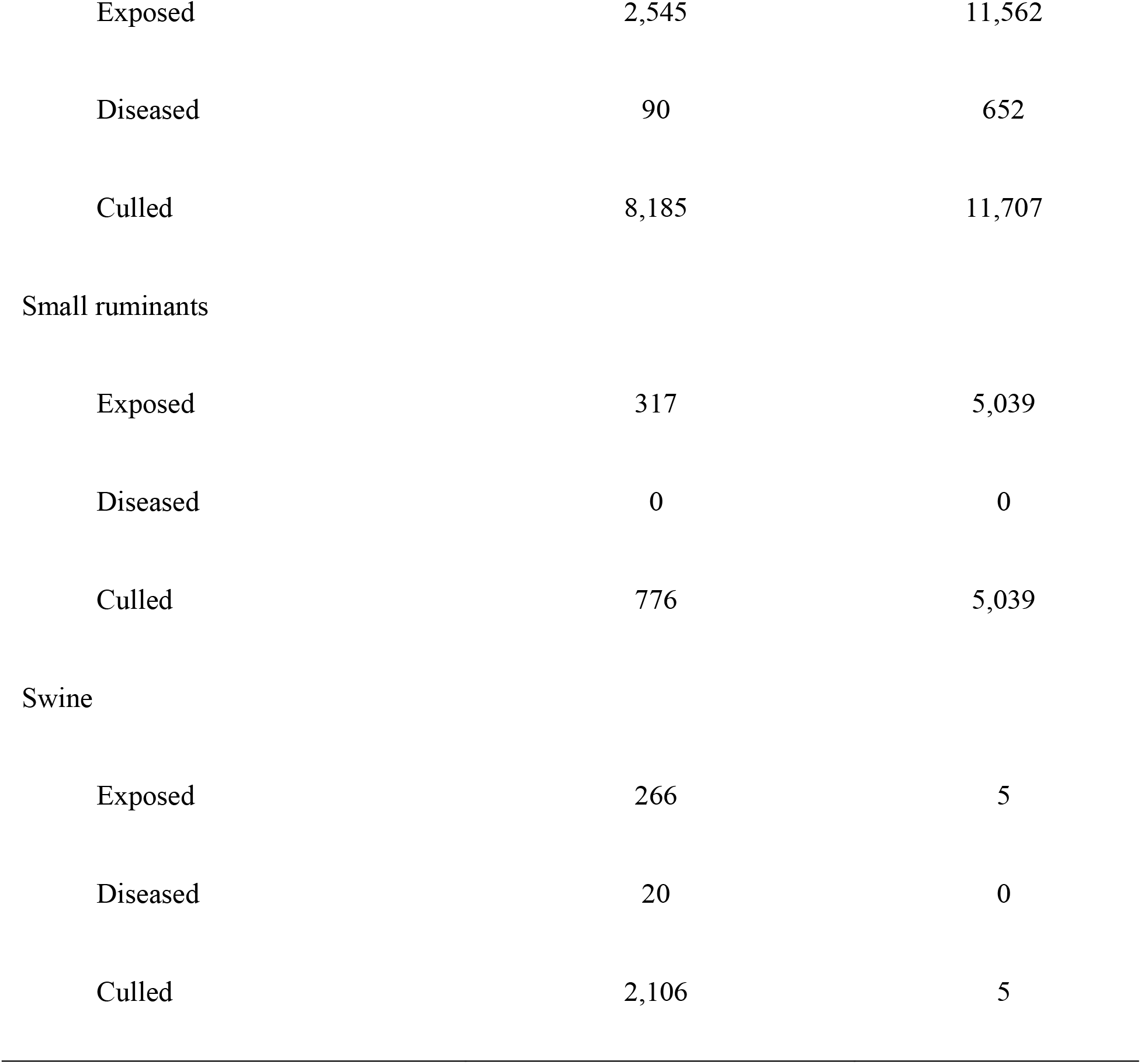
Years 2000 and 2001 outbreak descriptions are based on official veterinary state service archives, databases, and reports produced during and after the epidemic events (MAPA, 2002).

Control measures were implemented according to the current contingency plans published by the Brazilian Ministry of Agriculture, Livestock, and Supply (MAPA, 1993). It included establishing containment zones, with a 25 km radius measured from each outbreak farm, in which animals and animal product movement was restricted, and clinical inspection of susceptible animals was carried out routinely. Within the three km infected zones, susceptible animals on the infected farms and their immediate contact farms were culled, followed by disinfection of farms. Before repopulating farms, a 30-days quarantine was established with the introduction of unvaccinated naïve calves as sentinels, which were systematically submitted to clinical inspection and laboratory testing to discard possible persistence of the infectious viruses in the environment (MAPA, 2002). The 2000 outbreak occurred three months after the official withdrawal of FMD vaccination on May 1st (MAPA, 2000). The vaccination forbiddance was maintained in the state even after the outbreak started, as a strategy to keep the conditions required to obtain OIE free from FMD without vaccination zone status. The serosurveillance carried out as part of the activities to substantiate freedom from FMD involved 1,078 farms and 12,795 samples. The end of the health emergency state was declared in February 2001 and the final cost for the 2000 outbreak was estimated at US$ 3,7 million (MAPA, 2002).

Nevertheless, amid numerous FMD outbreaks being reported by Argentina and Uruguay animal health authorities (Correa Melo et al., 2002; Perez et al., 2004a, 2004b), on May 5th, 2001, a new introduction was identified in the municipality of Santana do Livramento, 5 km from the Uruguay border. Laboratory diagnosis confirmed a type A virus revealing the absence of epidemiological connection with the 2000 events (MAPA, 2002). The epidemic infected 30 farms in six municipalities, comprising a 27,053 km^**2**^ area, that can be divided into three distinct geographic areas: Santana do Livramento and three contiguous municipalities, Alegrete, Quaraí, and Dom Pedrito, and the municipality of Jari and Rio Grande, which are 225 km and 412 km distant from the index case, respectively. More information about the infected farms is described in Table 1. As an estimate of the delay between symptoms onset and official disease notification, as described earlier, the age of the oldest clinical lesion in days described for each outbreak was on average 5.03 with a standard deviation of 3.59 days. When we segregate only the outbreaks within the municipality of Rio Grande, where 18 farms were infected, the mean delay estimate rises to 6.94 (sd = 3.25), while for the remaining municipalities combined it drops to 2.17 (sd = 1.70)(MAPA, 2002).

In addition to the control measures outlined in the contingency plan implemented in the 2000 outbreak, for the 2001 outbreak, the authorities determined an immediate vaccination in cattle and buffaloes followed by revaccination after 30 to 40 days (MAPA, 2001). Serosurveillance was likewise conducted as part of the control measures but also to recover the status of free from FMD and resulted in almost 130,000 samples collected in 1,867 farms. The overall cost of the 2001 outbreak, including indemnities, was estimated at US$ 7,7 million, and the restrictions imposed to control the disease spread were only withdrawn in April 2002 (MAPA, 2002).

### 3.1 Estimated effective reproduction number

The median *R*_*t*_ estimated by the Bayesian model for the 2000 outbreak was 1.00 (90% CI 0.93, 1.10), while the rate of growth was negative 0.00051 (90% CI -0.012, 0.013). The model also estimated the median number of new daily cases for each infection date, the median was 0 with the 90% credible interval varying from 0 to 1 (Figure 2 A). For the 2001 outbreak, the median *R*_*t*_ was 1.60 (90% CI 1.50, 1.70), the median rate of growth was positive at 0.088 (90% CI 0.076, 0.100) and the median number of new daily cases was 3 (90% CI 3,4), Figure 2 B. Those parameters and their variations were also estimated throughout both outbreaks’ timelines (Figure 2).

**Figure 2:**
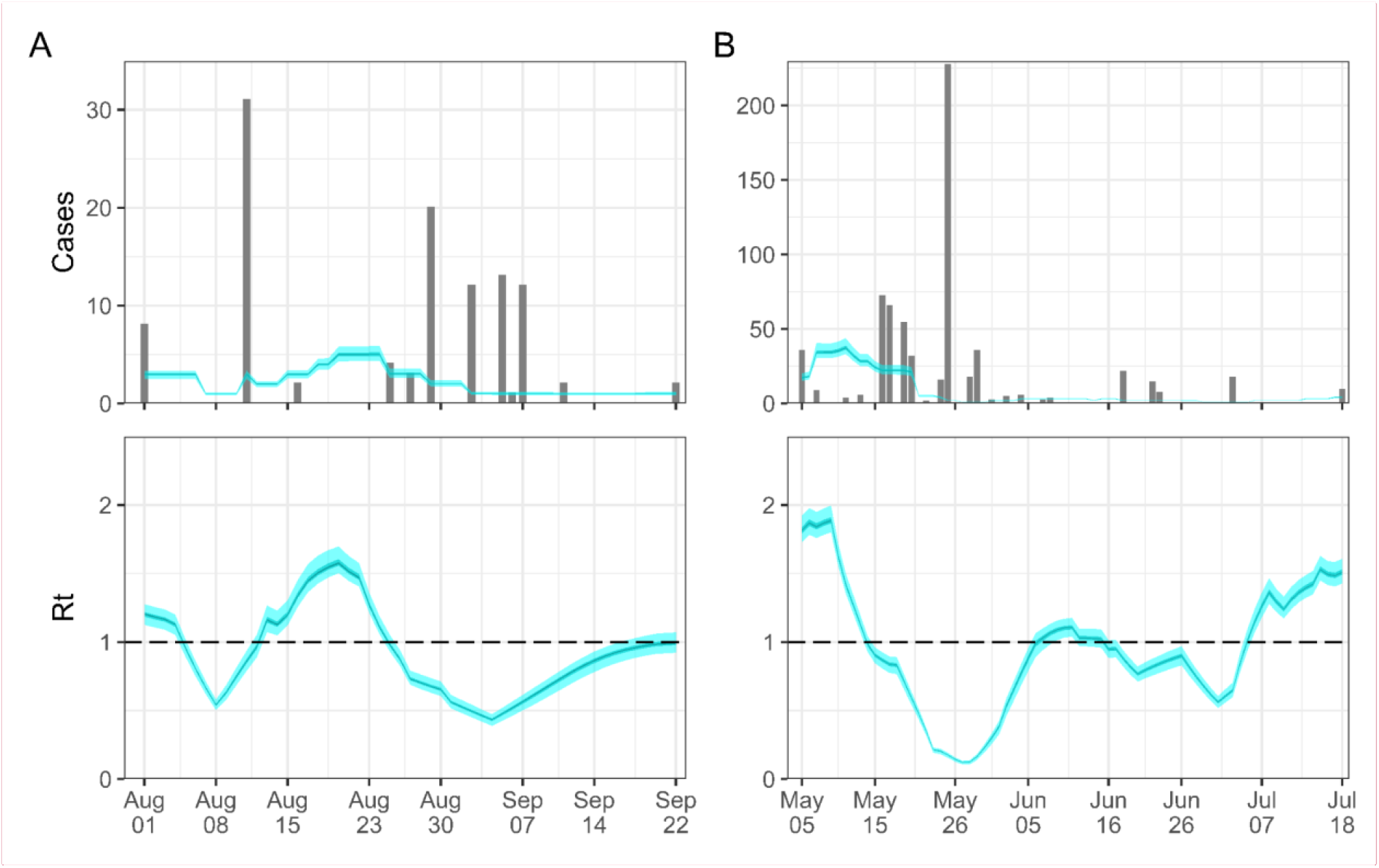
R_t_ estimates for the outbreaks in 2000 (A) and 2001 (B). For each estimate, the lightest blue ribbon illustrates 90% credible interval; the darker blue ribbon, 50% credible interval; and the darkest blue ribbon, 20% credible interval. Top panel: bars represent confirmed cases by date of notification and the ribbons illustrate estimated cases by date of infection. Bottom panel: time-varying estimate for the R_**t**_.

### 3.2 Static network analysis

We used the 2018 to 2020 movement network to reconstruct possible FMD epidemics involving the 2000 and 2001 infected farms. Results demonstrated that the total number of between-farm cattle movements was 692 batches adding up to a total of 5,246 animals, while in the 2001 infected network model there were 639 batches and 21,045 animals. The results of the cattle movement network analysis showed differences between the characteristics of the 2000 and 2001 infected network models (Table 2). The 2000 infected network had the largest number of nodes and edges, as well as the highest GWCC and centralization estimates. The GSCC was slightly higher in the 2001 network, which could suggest some directionality in the transmission. Furthermore, mean degree, diameter, and mean betweenness had similar estimations in both networks (Table 2).

**Table 2.** Cattle movement network model outcomes for the 2000 and 2001 outbreaks farms, based on between-farm cattle movement records from 2018 to 2020 (SEAPDR-RS, 2021).

|  | Foot-and-mouth disease outbreaks |  |
| --- | --- | --- |
|  | 2000 | 2001 |
| Mean degree | 1.99 | 2.07 |
| Nodes | 497 | 347 |
| Edges | 496 | 359 |
| Diameter | 3 | 4 |
| GSCC | 5 (1.01%) | 15 (4.32%) |
| GWCC | 463 (93.16%) | 173 (49.86%) |
| Mean betweenness | 21.95 | 21.19 |
| Centralization | 0.46 | 0.27 |

The permutation-based approach showed that the distribution patterns of outbreak farms within both movement networks were not significant. This result suggests that assuming the same infected farms for the 2000 and 2001 outbreaks modeled independently in the 2018 to 2020 network, the cattle movement alone would not explain transmission pathways between farms.

## 4. Discussion

The reassessment of the 2000 and 2001 FMD reintroductions in the state of Rio Grande do Sul allowed the description of the characteristics of the outbreaks with important lessons learned from two distinct epidemiological events. More importantly, the data collected was used to estimate FMD spread parameters, clearly demonstrating two different epidemics, with 2000 being more limited (median *R*_*t*_ around 1) when compared to 2001 (median *R*_*t*_ above 1). We also used the 2018 to 2020 cattle movement network as a standard to benchmark transmission dynamics of infected farms independently for the 2000 and 2001 outbreaks. We found similarities in the characteristics of the simulated transmission pattern between farms and the reported results of the official outbreak investigation.

Investigation reports indicated that the illegal import of cattle infected with a type O FMD virus from the northern region of Argentina to a farm in the municipality of Jóia, 145 km distant from the border, was the origin of the 2000 outbreak (MAPA, 2002). Later studies demonstrated a close genetic relationship between the Argentina virus strain and the ones isolated in Uruguay and Brazil during the same year, suggesting that the transborder movement played an important role in the virus dissemination (Malirat et al., 2007; Mattion et al., 2004). The type A virus isolated in the 2001 outbreak was also genetically matched at that time with the strains isolated in Uruguay and Argentina (MAPA, 2002). These findings were corroborated by other genetics studies carried out after the events (König et al., 2007; Mattion et al., 2004).

The 2000 outbreak was limited to 22 infected farms in four municipalities which could be attributed to a concentration of low animal density farms, an intense movement of milk tank trucks, and unofficial animal movements between those farms (MAPA, 2002). The low-density area combined with the epidemics countermeasures adopted, and the herd residual immunity due to recent vaccination withdrawal three months before the first case was identified (MAPA, 2000), might explain both the low *R*_*t*_ and rate of growth throughout the epidemic. This limited spread is similar to outbreaks in historically endemic regions like Thailand, where a study found a subdistrict reproduction number (*R*_*sd*_) ranging from 1.04 and 1.07 (Arjkumpa et al., 2021). However, a year had passed since the last vaccination when FMD was reintroduced in 2001, thus animal population immunity was probably lower than in 2000. According to the official outbreak investigation reports, the more intense between-farm movement within municipalities and delayed notification of identified cases by owners, especially in the farms located in the municipality of Rio Grande, were the main factors that contributed to the spread of the disease (MAPA, 2002). We demonstrated that *R*_*t*_ in the 2001 outbreak was 60% greater than the 2000 outbreak and substantially above one, indicating a sustained increase in disease transmission (Anderson & May, 1995). Other studies estimated the trend on the FMD epidemic curve using the herd reproduction ratio (*R*_*h*_) and showed significantly higher results. In 2001, the outbreak in Argentina had a mean *R*_*h*_ of 2.4 (Perez et al., 2004a) and in The Netherlands, the mean estimate for the period before the first disease notification was *R*_*h*_= 2.6, and after the countermeasures actions were implemented it dropped to *R*_*h*_= 0.71 (Bouma et al., 2003). In the 2004 Peru outbreak, *R*_*h*_was 5.3 at the beginning of the epidemic and declined to 1.3 at the end of the outbreak (Estrada et al., 2008). Studies in endemic regions, such as Ethiopia, using the basic reproduction number estimated the average *R*_0_ of 1.68 in the smallholder production systems and 1.98 in commercial dairy farms (Tadesse et al., 2019). In addition, we observed a sharp decrease two weeks after the beginning of the 2001 outbreak (Figure 2) which could be explained by the mass vaccination implemented four days after the first case was identified and reported (MAPA, 2001) combined with other countermeasures described earlier in this study. The effect of vaccination on the decline of the FMD epidemic curve is also demonstrated in other field investigations (Estrada et al., 2008; Perez et al., 2004a) and experimental studies (Orsel et al., 2005).

The main hypothesis about the dissemination of the 2000 outbreak from Jóia to other municipalities was attributed to unofficial cattle movement and between-farm indirect contact through artificial insemination technical assistance (MAPA, 2002). Investigations also assessed retrospectively the registries of susceptible animal movements for the past 90 days before the start of the outbreak and followed investigations in contact properties. It concluded that the formal animal movement was not accountable for the spread of the disease and reinforced the assumption that the outbreaks remained limited to the control zone. In the 2001 outbreaks, no direct epidemiological relationships between the six municipalities with infected farms were identified in the field investigation, which included animal movement traceback. The main hypothesis for the 2001 outbreak was that all six outbreaks were independent, and originated directly from Uruguay. In the outbreak in the municipality of Jarí, the furthest from the border among the affected municipalities, direct relationships were found between the affected farm owner and animals raised in Uruguay, with demonstrated direct movement of cattle from Uruguay (MAPA, 2002). Assuming the same infected farms, independently for the 2000 and 2001 outbreaks, using the 2018 to 2020 movement network as standard, the results of the permutation-based approach (k-statistic) reiterate the epidemic hypotheses that the cattle movement did not explain transmission pathways between farms. Also, once comprehended the time gap between the outbreak events and the data used in this study, the analysis of the infected network can capture complex population-level structure and connectivity, in order to estimate the extent of disease spread in some simulated scenarios. The results demonstrate mainly similarities between infected networks, except for the 2000 network which has higher values for nodes and edges counts, as well as for GWCC and centralization estimates. Thus, considering the assumptions outlined above, the present network analysis suggests that, had the reintroduction happened in a population with a movement pattern like the one modeled in this study for the 2000 infected network, a greater number of farms would be involved in the movement network but with low connectivity between them. We also would expect farms that acted as hubs, which receive animals from many other farms, hence requiring higher attention when planning the countermeasures of an outbreak (Chaters et al., 2019; Dubé et al., 2009; Wasserman & Faust, 1994).

## 5. Limitation and further remarks

The information associated with the FMD reintroductions in the State of Rio Grande do Sul was limited to farm-level demographics information at that time, however, because of limited technical information about the virus genetics was not captured. Another important limitation was the lack of electronic data on the between-farm movements, which limited our ability to reconstruct those dissemination events with real movement data.

The lack of information, associated with the small 2000 FMD epidemic, narrowed the availability of models to estimate transmission parameters, requiring a pragmatic choice of methods. In the analysis of the *R*_*t*_ results, direct comparisons between different outbreaks’ reproduction numbers have major restrictions due to several causes, such as divergences in statistical methods, inherent models limitations, including the ones used in this paper (Abbott et al., 2020), differences in models assumptions, and how it deals with uncertainties derived from the infection dynamics and data biases (Delamater et al., 2019; van Andel et al., 2021; Vegvari et al., 2021). Furthermore, the variation of virulence and infectivity between different FMD virus strains, and heterogeneity of the host animal species may also affect the transmission dynamics (Bravo De Rueda et al., 2015; Kitching et al., 2006).

In addition to the previously exposed limitation arriving from the gap between the outbreaks events and the data used in the analysis of the movement network, the use of static networks by itself should be employed with caution due to limitations associated with animal movement aggregation and network stability (Lentz et al., 2016; Machado et al., 2021). Also, the present network analysis did not account for other host species movements, nor did it incorporate spatial information, which is highly related to FMD transmission, thus suggesting the demands for future work.

Once addressed the limitations outlined throughout this paper, the models used in this study may be useful choices for assessing the transmission dynamics and support control measures in the future.

## 6. Conclusion

We revised historical FMD outbreaks in the state of Rio Grande do Sul, Brazil, in 2000 and 2001. For 2000 outbreaks the median *R*_*t*_ was estimated around 1, which showed consistency with the limited epidemic described in the investigation reports, while for the 2001 outbreak, the median *R*_*t*_ was significantly above 1, therefore corroborating with the wider spread of the disease observed. The results could be practically interpreted by the differences in the serotypes of the virus involved, the density of the animal population, delayed notification of identified cases, and the unequal levels of immunity considering the longest interval since the last vaccination.

Additionally, we modeled the contact networks using the 2018 to 2020 cattle movement network as standard and assuming the same infected farms independently for the 2000 and 2001 outbreaks. We found similarities in the characteristics of the simulated transmission pattern between farms and the reported results of the official outbreak investigation. The models and results presented in this study may be useful for assessing the transmission dynamics and support control measures in the future.

## Acknowledgments

This work was funded by the Fundo de Desenvolvimento e Defesa Sanitária Animal (FUNDESA-RS).

## Authors’ contributions

JMNC and GM conceived the study. JMNC, NCC, and GM participated in the design of the study. FHSG participated in data collection and validation. JMNC conducted data processing and cleaning and designed the model with the assistance of GM. JMNC, NCC, and GM designed the computational analysis. JMNC and NCC conducted the formal coding. GM and JMNC wrote and edited the manuscript. All authors discussed the results and critically reviewed the manuscript. GM secured the funding.

## Ethical statement

The authors confirm the ethical policies of the journal, as noted on the journal’s author guidelines page. Since this work did not involve animal sampling nor questionnaire data collection by the researchers, there was no need for ethics permits.

## Data Availability Statement

The data that support the findings of this study are not publicly available and are protected by confidential agreements, therefore, are not available.

## References

Abbott, S., Hellewell, J., Thompson, R. N., Sherratt, K., Gibbs, H. P., Bosse, N. I., Munday, J. D., Meakin, S., Doughty, E. L., Chun, J. Y., Chan, Y.-W. D., Finger, F., Campbell, P., Endo, A., Pearson, C. A. B., Gimma, A., Russell, T., Flasche, S., Kucharski, A. J., … Funk, S. (2020). Estimating the time-varying reproduction number of SARS-CoV-2 using national and subnational case counts. Wellcome Open Research, 5, 112. https://doi.org/10.12688/wellcomeopenres.16006.2

Abbott, S., JoeHickson, Hamada S. Badr, Funk, S., Monticone, P., Ellis, P., Jdmunday Allen, J., Pearson, C. A. B., DeWitt, M., Bosse, N., & Meakin, S. (2021). epiforecasts/EpiNow2: Beta release (1.3.3) [Computer software]. Zenodo. https://doi.org/10.5281/ZENODO.3957489

Anderson, R. M., & May, R. M. (1995). Infectious diseases of humans: Dynamics and control (Reprinted). Oxford university press.

Arjkumpa, O., Picasso-Risso, C., Perez, A., & Punyapornwithaya, V. (2021). Subdistrict-Level Reproductive Number for Foot and Mouth Disease in Cattle in Northern Thailand. Frontiers in Veterinary Science, 8(November), 5–7. https://doi.org/10.3389/fvets.2021.757132

Arzt, J., Baxt, B., Grubman, M. J., Jackson, T., Juleff, N., Rhyan, J., Rieder, E., Waters, R., & Rodriguez, L. L. (2011). The Pathogenesis of Foot-and-Mouth Disease II: Viral Pathways in Swine, Small Ruminants, and Wildlife; Myotropism, Chronic Syndromes, and Molecular Virus-Host Interactions. Transboundary and Emerging Diseases, 58(4), 305–326. https://doi.org/10.1111/j.1865-1682.2011.01236.x

Arzt, J., Juleff, N., Zhang, Z., & Rodriguez, L. L. (2011). The Pathogenesis of Foot-and-Mouth Disease I: Viral Pathways in Cattle. Transboundary and Emerging Diseases, 58(4), 291–304. https://doi.org/10.1111/J.1865-1682.2011.01204.X

Bouma, A., Elbers, A. R. W., Dekker, A., Koeijer, A. D., Bartels, C., Vellema, P., Wal, P. V. D., Rooij, E. M. A. V., Pluimers, F. H., & Jong, M. C. M. D. (2003). The foot-and-mouth disease epidemic in The Netherlands in 2001. Preventive Veterinary Medicine, 57(3), 155–166. https://doi.org/10.1016/S0167-5877(02)00217-9

Bravo De Rueda, C., De Jong, M. C., Eblé, P. L., & Dekker, A. (2015). Quantification of transmission of foot-and-mouth disease virus caused by an environment contaminated with secretions and excretions from infected calves. Veterinary Research, 46(1). https://doi.org/10.1186/s13567-015-0156-5

Cabezas, A. H., Sanderson, M. W., & Volkova, V. V. (2021). Modeling Intervention Scenarios During Potential Foot-and-Mouth Disease Outbreaks Within U.S. Beef Feedlots. Frontiers in Veterinary Science, 8(February), 1–14. https://doi.org/10.3389/fvets.2021.559785

Cardenas, N. C., Sykes, A. L., Lopes, F. P. N., & Machado, G. (2022). Multiple species animal movements: Network properties, disease dynamics and the impact of targeted control actions. Veterinary Research, 53(1), 1–16. https://doi.org/10.1186/s13567-022-01031-2

Cardenas, N. C., VanderWaal, K., Veloso, F. P., Galvis, J. O. A., Amaku, M., & Grisi-Filho, J. H. H. (2021). Spatio-temporal network analysis of pig trade to inform the design of risk-based disease surveillance. Preventive Veterinary Medicine, 189(February), 105314. https://doi.org/10.1016/j.prevetmed.2021.105314

Chaters, G. L., Johnson, P. C. D., Cleaveland, S., Crispell, J., Glanville, W. A. D., Doherty, T., Matthews, L., Mohr, S., Nyasebwa, O. M., Rossi, G., Salvador, L. C. M., Swai, E., & Kao, R. R. (2019). Analysing livestock network data for infectious disease control: An argument for routine data collection in emerging economies. Philosophical Transactions of the Royal Society B: Biological Sciences, 374(1776). https://doi.org/10.1098/rstb.2018.0264

Colenutt, C., Brown, E., Nelson, N., Paton, D. J., Eblé, P., Dekker, A., Gonzales, J. L., & Gubbins, S. (2020). Quantifying the transmission of foot-and-mouth disease virus in cattle via a contaminated environment. MBio, 11(4), 1–13. https://doi.org/10.1128/mBio.00381-20

Cori, A., Ferguson, N. M., Fraser, C., & Cauchemez, S. (2013). A new framework and software to estimate time-varying reproduction numbers during epidemics. American Journal of Epidemiology, 178(9), 1505–1512. https://doi.org/10.1093/aje/kwt133

Correa Melo, E., Saraiva, V. E. V., & Astudillo, V. M. (2002). Review of the status of foot and mouth disease in countries of South America and approaches to control and eradication: -EN--FR--ES-. Revue Scientifique et Technique de l’OIE, 21(3), 429–436. https://doi.org/10.20506/rst.21.3.1350

Csardi, G & Nepusz, T. (2006). The igraph software package for complex network research. In InterJournal: Vol. Complex Systems (p. 1695). https://igraph.org

Delamater, P. L., Street, E. J., Leslie, T. F., Yang, Y. T., & Jacobsen, K. H. (2019). Complexity of the Basic Reproduction Number (R _0_). Emerging Infectious Diseases, 25(1), 1–4. https://doi.org/10.3201/eid2501.171901

Dietz, K. (1993). The estimation of the basic reproduction number for infectious diseases. Statistical Methods in Medical Research, 2(1), 23–41. https://doi.org/10.1177/096228029300200103

Dubé, C., Ribble, C., Kelton, D., & McNab, B. (2009). A Review of Network Analysis Terminology and its Application to Foot-and-Mouth Disease Modelling and Policy Development. Transboundary and Emerging Diseases, 56(3), 73–85. https://doi.org/10.1111/j.1865-1682.2008.01064.x

Estrada, C., Perez, A. M., & Turmond, M. C. (2008). Herd Reproduction Ratio and Time-Space Analysis of a Foot-and-mouth Disease Epidemic in Peru in 2004. Transboundary and Emerging Diseases, 55(7), 284–292. https://doi.org/10.1111/j.1865-1682.2008.01023.x

EUFMD. (2020). A field guide to estimating the age of Foot-and-Mouth disease lesions. European Commission for the Control of Foot-and-Mouth Disease. https://eufmdlearning.works/mod/page/view.php?id=3675

FAO. (2018). Global Foot-and-Mouth Disease Situation—October 2018. In Foot-and-Mouth Disease Situation—Monthly Report (Vol. 10, p. 32). Food and Agriculture Organization of the United Nations. http://www.fao.org/fileadmin/user_upload/eufmd_new/docs/October_GMR_2018.pdf

Ferguson, N. M., Donnelly, C. A., & Anderson, R. M. (2001). The foot-and-mouth epidemic in Great Britain: Pattern of spread and impact of interventions. Science, 292(5519), 1155–1160. https://doi.org/10.1126/science.1061020

Fèvre, E. M., Bronsvoort, B. M. D. C., Hamilton, K. A., & Cleaveland, S. (2006). Animal movements and the spread of infectious diseases. Trends in Microbiology, 14(3), 125–131. https://doi.org/10.1016/j.tim.2006.01.004

Firestone, S. M., Hayama, Y., Lau, M. S. Y., Yamamoto, T., Nishi, T., Bradhurst, R. A., Demirhan, H., Stevenson, M. A., & Tsutsui, T. (2020). Transmission network reconstruction for foot-and-mouth disease outbreaks incorporating farm-level covariates. PLoS ONE, 15(7 July), 1– 17. https://doi.org/10.1371/journal.pone.0235660

Gomez, F., Prieto, J., Galvis, J., Moreno, F., & Vargas, J. (2019). Identification of Super-Spreaders of Foot-And-Mouth Disease in the cattle transportation network: The 2018 outbreak case in Cesar (Colombia). Proceedings of 2019 IEEE World Conference on Complex Systems, WCCS 2019, 5–10. https://doi.org/10.1109/ICoCS.2019.8930765

Gonçalves, V. S. P., & de Moraes, G. M. (2017). The application of epidemiology in national veterinary services: Challenges and threats in Brazil. Preventive Veterinary Medicine, 137, 140–146. https://doi.org/10.1016/j.prevetmed.2016.11.018

Gostic, K. M., McGough, L., Baskerville, E. B., Abbott, S., Joshi, K., Tedijanto, C., Kahn, R., Niehus, R., Hay, J. A., Salazar, P. M. D., Hellewell, J., Meakin, S., Munday, J. D., Bosse, N. I., Sherrat, K., Thompson, R. N., White, L. F., Huisman, J. S., Scire, J., … Cobey, S. (2020). Practical considerations for measuring the effective reproductive number, Rt. PLoS Computational Biology, 16(12), 1–21. https://doi.org/10.1371/journal.pcbi.1008409

Grubman, M. J., & Baxt, B. (2004). Foot-and-mouth disease. Clinical Microbiology Reviews, 17(2), 465–493. https://doi.org/10.1128/cmr.17.2.465-493.2004

Hayama, Y., Firestone, S. M., Stevenson, M. A., Yamamoto, T., Nishi, T., Shimizu, Y., & Tsutsui, T. (2019). Reconstructing a transmission network and identifying risk factors of secondary transmissions in the 2010 foot-and-mouth disease outbreak in Japan. Transboundary and Emerging Diseases, 66(5), 2074–2086. https://doi.org/10.1111/tbed.13256

Haydon, D. T., Chase-Topping, M., Shaw, D. J., Matthews, L., Friar, J. K., Wilesmith, J., & Woolhouse, M. E. J. (2003). The construction and analysis of epidemic trees with reference to the 2001 UK foot-and-mouth outbreak. Proceedings of the Royal Society B: Biological Sciences, 270(1511), 121–127. https://doi.org/10.1098/rspb.2002.2191

Jombart, T., Cori, A., Didelot, X., Cauchemez, S., Fraser, C., & Ferguson, N. (2014). Bayesian Reconstruction of Disease Outbreaks by Combining Epidemiologic and Genomic Data. PLoS Computational Biology, 10(1). https://doi.org/10.1371/journal.pcbi.1003457

Junker, F., Ilicic-Komorowska, J., Tongeren, F. V., & Junker, F. (2009). Impact of Animal Disease Outbreaks and Alternative Control Practices on Agricultural Markets and Trade THE CASE OF FMD. https://doi.org/10.1787/221275827814

Keeling, M. J. (2005). Models of foot-and-mouth disease. Proceedings of the Royal Society B: Biological Sciences, 272(1569), 1195–1202. https://doi.org/10.1098/rspb.2004.3046

Kitching, R. P., & Alexandersen, S. (2002). Clinical variation in foot and mouth disease: Cattle. OIE Revue Scientifique et Technique, 21(3), 499–504. https://doi.org/10.20506/rst.21.3.1367

Kitching, R. P., Thrusfield, M. V., & Taylor, N. M. (2006). Use and abuse of mathematical models: An illustration from the 2001 foot and mouth disease epidemic in the United Kingdom. OIE Revue Scientifique et Technique, 25(1), 293–311. https://doi.org/10.20506/rst.25.1.1665

Knight-Jones, T. J. D., Robinson, L., Charleston, B., Rodriguez, L. L., Gay, C. G., Sumption, K. J., & Vosloo, W. (2016). Global Foot-and-Mouth Disease Research Update and Gap Analysis: 2. Transboundary and Emerging Diseases, 63(S1), 3–13. https://doi.org/10.1111/tbed.12528

Knight-Jones, T. J. D., & Rushton, J. (2013). The economic impacts of foot and mouth disease—What are they, how big are they and where do they occur? Preventive Veterinary Medicine, 112(3– 4), 162–173. https://doi.org/10.1016/j.prevetmed.2013.07.013

König, G. A., Palma, E. L., Maradei, E., & Piccone, M. E. (2007). Molecular epidemiology of foot-and-mouth disease virus types A and O isolated in Argentina during the 2000-2002 epizootic. Veterinary Microbiology, 124(1–2), 1–15. https://doi.org/10.1016/j.vetmic.2007.03.015

Lentz, H. H. K., Koher, A., Hövel, P., Gethmann, J., Sauter-Louis, C., Selhorst, T., & Conraths, F. J. (2016). Disease spread through animal movements: A static and temporal network analysis of pig trade in Germany. PLoS ONE, 11(5), 1–32. https://doi.org/10.1371/journal.pone.0155196

Lyra, T. M. P., & Silva, J. A. (2004). A febre aftosa no Brasil, 1960-2002. Arquivo Brasileiro de Medicina Veterinária e Zootecnia, 56(5), 565–576. https://doi.org/10.1590/s0102-09352004000500001

Machado, G., Galvis, J. A., Lopes, F. P. N., Voges, J., Medeiros, A. A. R., & Cárdenas, N. C. (2021). Quantifying the dynamics of pig movements improves targeted disease surveillance and control plans. Transboundary and Emerging Diseases, 68(3), 1663–1675. https://doi.org/10.1111/tbed.13841

Malirat, V., Barros J. J. F. de, Bergmann, I. E., Campos, R. de M., Neitzert, E., Costa E. V. da, Silv E. E. da, Falczuk, A. J., Pinheiro, D. S. B., Vergara N. de, Cirvera, J. L. Q., Maradei, E., & Landro, R. D. (2007). Phylogenetic analysis of foot-and-mouth disease virus type O re-emerging in free areas of South America. Virus Research, 124(1–2), 22–28. https://doi.org/10.1016/j.virusres.2006.09.006

MAPA. (1993). PORTARIA no 121, DE 29 DE MARÇO DE 1993. Ministério Da Agricultura, Pecuária e Abastecimento. https://sistemasweb.agricultura.gov.br/sislegis/action/detalhaAto.do?method=abreLegislacaoFederal&chave=50674&tipoLegis=A

MAPA. (2000). INSTRUÇÃO NORMATIVA No 13, DE 19 DE MAIO DE 2000. Ministério Da Agricultura, Pecuária e Abastecimento. https://sistemasweb.agricultura.gov.br/sislegis/action/detalhaAto.do?method=abreLegislacaoFederal&chave=50674&tipoLegis=A

MAPA. (2001). INSTRUÇÃO NORMATIVA N° 11, DE 9 DE MAIO DE 2001. Ministério Da Agricultura, Pecuária e Abastecimento. https://sistemasweb.agricultura.gov.br/sislegis/action/detalhaAto.do?method=abreLegislacaoFederal&chave=50674&tipoLegis=A

MAPA. (2002). Eliminação dos focos de febre aftosa no Estado do Rio Grande do Sul. Ministério da Agricultura, Pecuária e Abastecimento.

Mardones, F., Perez, A., Sanchez, J., Alkhamis, M., & Carpenter, T. (2010). Parameterization of the duration of infection stages of serotype O foot-and-mouth disease virus: An analytical review and meta-analysis with application to simulation models. Veterinary Research, 41(4). https://doi.org/10.1051/vetres/2010017

Mattion, N., König, G., Seki, C., Smitsaart, E., Maradei, E., Robiolo, B., Duffy, S., León, E., Piccone, M., Sadir, A., Bottini, R., Cosentino, B., Falczuk, A., Maresca, R., Periolo, O., Bellinzoni, R., Espinoza, A., Torre, J. L., & Palma, E. L. (2004). Reintroduction of foot-and-mouth disease in Argentina: Characterisation of the isolates and development of tools for the control and eradication of the disease. Vaccine, 22(31–32), 4149–4162. https://doi.org/10.1016/j.vaccine.2004.06.040

Mayen, F. L. (2003). Foot and mouth disease in Brazil and its control—An overview of its history, present situation and perspectives for eradication. Veterinary Research Communications, 27(2), 137–148. https://doi.org/10.1023/A:1022863221356

Merl, D., Johnson, L. R., Gramacy, R. B., & Mangel, M. (2009). A statistical framework for the adaptive management of epidemiological interventions. PLoS ONE, 4(6). https://doi.org/10.1371/journal.pone.0005807

Mielke, S. R., & Garabed, R. (2020). Environmental persistence of foot-and-mouth disease virus applied to endemic regions. Transboundary and Emerging Diseases, 67(2), 543–554. https://doi.org/10.1111/TBED.13383

Miller, M., Liu, L., Shwiff, S., & Shwiff, S. (2019). Macroeconomic impact of foot-and-mouth disease vaccination strategies for an outbreak in the Midwestern United States: A computable general equilibrium. Transboundary and Emerging Diseases, 66(1), 156–165. https://doi.org/10.1111/tbed.12995

Muroga, N., Hayama, Y., Yamamoto, T., Kurogi, A., Tsuda, T., & Tsutsui, T. (2012). The 2010 foot-and-mouth disease epidemic in Japan. Journal of Veterinary Medical Science, 74(4), 399– 404. https://doi.org/10.1292/jvms.11-0271

Musser, J. M. B. (2004). A practitioner’s primer on foot-and-mouth disease. Journal of the American Veterinary Medical Association, 224(8), 1261–1268. https://doi.org/10.2460/javma.2004.224.1261

Nakajo, K., & Nishiura, H. (2022). Estimation of R(t) based on illness onset data: An analysis of 1907–1908 smallpox epidemic in Tokyo. Epidemics, 38. https://doi.org/10.1016/j.epidem.2022.100545

Naranjo, J., & Cosivi, O. (2013). Elimination of foot-and-mouth disease in South America: Lessons and challenges. Philosophical Transactions of the Royal Society B: Biological Sciences, 368(1623). https://doi.org/10.1098/rstb.2012.0381

OIE. (2021). Recognition of the Foot and Mouth Disease Status of Members (p. 4). World Organisation for Animal Health. https://www.oie.int/app/uploads/2021/05/a-r13-2021-fmd.pdf

Orsel, K., Dekker, A., Bouma, A., Stegeman, J. A., & De Jong, M. C. M. (2005). Vaccination against foot and mouth disease reduces virus transmission in groups of calves. Vaccine, 23(41), 4887–4894. https://doi.org/10.1016/j.vaccine.2005.05.014

PANAFTOSA. (2020). Informe de situacion de los Programas de Erradicación de la Febre Aftosa en Sudamérica y Panamá en 2019. Centro Panamericano de Fiebre Aftosa. https://www.paho.org/es/documentos/informe-situacion-programas-erradicacion-fiebre-aftosa-sudamerica-panama-ano-2019

Paton, D. J., Gubbins, S., & King, D. P. (2018). Understanding the transmission of foot-and-mouth disease virus at different scales. Current Opinion in Virology, 28(Figure 2), 85–91. https://doi.org/10.1016/j.coviro.2017.11.013

Pendell, D. L., Leatherman, J., Schroeder, T. C., & Alward, G. S. (2007). The Economic Impacts of a Foot-And-Mouth Disease Outbreak: A Regional Analysis. Journal of Agricultural and Applied Economics, 39(1), 19–33. https://doi.org/10.1017/S1074070800028911

Perez, A. M., Ward, M. P., & Carpenter, T. E. (2004a). Control of a foot-and-mouth disease epidemic in Argentina. Preventive Veterinary Medicine, 65(3–4), 217–226. https://doi.org/10.1016/j.prevetmed.2004.08.002

Perez, A. M., Ward, M. P., & Carpenter, T. E. (2004b). Epidemiological investigations of the 2001 foot-and-mouth disease outbreak in Argentina. Veterinary Record, 154(25), 777–782. https://doi.org/10.1136/vr.154.25.777

Pomeroy, L. W., Bansal, S., Tildesley, M., Moreno-Torres, K. I., Moritz, M., Xiao, N., Carpenter, T. E., & Garabed, R. B. (2017). Data-Driven Models of Foot-and-Mouth Disease Dynamics: A Review. Transboundary and Emerging Diseases, 64(3), 716–728. https://doi.org/10.1111/tbed.12437

Probert, W. J. M., Jewell, C. P., Werkman, M., Fonnesbeck, C. J., Goto, Y., Runge, M. C., Sekiguchi, S., Shea, K., Keeling, M. J., Ferrari, M. J., & Tildesley, M. J. (2018). Real-time decision-making during emergency disease outbreaks. PLoS Computational Biology, 14(7). https://doi.org/10.1371/journal.pcbi.1006202

R Core Team. (2019). R: A language and environment for statistical computing. R Foundation for Statistical Computing. https://www.R-project.org/

Rueda, C. B. D., Jong, M. C. D., Eblé, P. L., & Dekker, A. (2015). Quantification of transmission of foot-and-mouth disease virus caused by an environment contaminated with secretions and excretions from infected calves. Veterinary Research, 46(1). https://doi.org/10.1186/s13567-015-0156-5

Ruget, A. S., Rossi, G., Pepler, P. T., Beaunée, G., Banks, C. J., Enright, J., & Kao, R. R. (2021). Multi-species temporal network of livestock movements for disease spread. Applied Network Science, 6(1). https://doi.org/10.1007/s41109-021-00354-x

SEAPDR-RS. (2021). Secretaria da Agricultura, Pecuária e Desenvolvimento Rural do Rio Grande do Sul. https://www.agricultura.rs.gov.br/vigilancia-defesa-sanitaria-animal

Sherratt, K., Abbott, S., Meakin, S. R., Hellewell, J., Munday, J. D., Bosse, N., Jit, M., & Funk, S. (2021). Exploring surveillance data biases when estimating the reproduction number: With insights into subpopulation transmission of COVID-19 in England. Philosophical Transactions of the Royal Society B: Biological Sciences, 376(1829). https://doi.org/10.1098/rstb.2020.0283

Tadesse, B., Molla, W., Mengsitu, A., & Jemberu, W. T. (2019). Transmission dynamics of foot and mouth disease in selected outbreak areas of northwest Ethiopia. Epidemiology and Infection, 147. https://doi.org/10.1017/S0950268819000803

Tildesley, M. J., & Keeling, M. J. (2009). Is R 0 a good predictor of final epidemic size: Foot-and-mouth disease in the UK. Journal of Theoretical Biology, 258(4), 623–629. https://doi.org/10.1016/j.jtbi.2009.02.019

van Andel, M., Tildesley, M. J., & Gates, M. C. (2021). Challenges and opportunities for using national animal datasets to support foot-and-mouth disease control. In Transboundary and Emerging Diseases (Vol. 68, Issue 4). https://doi.org/10.1111/tbed.13858

VanderWaal, K. L., Picasso, C., Enns, Eva. A., Craft, M. E., Alvarez, J., Fernandez, F., Gil, A., Perez, A., & Wells, S. (2016). Network analysis of cattle movements in Uruguay: Quantifying heterogeneity for risk-based disease surveillance and control. Preventive Veterinary Medicine, 123(6), 12–22. https://doi.org/10.1016/j.prevetmed.2015.12.003

Vegvari, C., Abbott, S., Ball, F., Brooks-Pollock, E., Challen, R., Collyer, B. S., Dangerfield, C., Gog, J. R., Gostic, K. M., Heffernan, J. M., Hollingsworth, T. D., Isham, V., Kenah, E., Mollison, D., Panovska-Griffiths, J., Pellis, L., Roberts, M. G., Scalia Tomba, G., Thompson, R. N., & Trapman, P. (2021). Commentary on the use of the reproduction number R during the COVID-19 pandemic. Statistical Methods in Medical Research, 1–11. https://doi.org/10.1177/09622802211037079

Wasserman, S., & Faust, K. (1994). Social Network Analysis: Methods and Applications (1st ed.). Cambridge University Press. https://doi.org/10.1017/CBO9780511815478

